# Intense bitterness of molecules: machine learning for expediting drug discovery

**DOI:** 10.1101/2020.06.24.168914

**Authors:** Eitan Margulis, Ayana Dagan-Wiener, Robert S. Ives, Sara Jaffari, Karsten Siems, Masha Y. Niv

## Abstract

Drug development is a long, expensive and multistage process geared to achieving safe drugs with high efficacy. A crucial prerequisite for completing the medication regimen for oral drugs, particularly for pediatric and geriatric populations, is achieving taste that does not hinder compliance. Currently, the aversive taste of drugs is tested in late stages of clinical trials. This can result in the need to reformulate, potentially resulting in the use of more animals for additional toxicity trials, increased financial costs and a delay in release to the market. Here we present BitterIntense, a machine learning tool that classifies molecules into “very bitter” or “not very bitter”, based on their chemical structure. The model, trained on chemically diverse compounds, has above 80% accuracy on several test sets. BitterIntense suggests that intense bitterness does not correlate with toxicity and hepatotoxicity of drugs and that the prevalence of very bitter compounds among drugs is lower than among microbial compounds. BitterIntense allows quick and easy prediction of strong bitterness of compounds of interest for food and pharma industries. We estimate that implementation of BitterIntense or similar tools early in drug discovery and development process may lead to reduction in delays, in animal use and in overall financial burden.

**Significance Statement:** Drug development integrates increasingly sophisticated technologies, but extreme bitterness of drugs remains a poorly addressed cause of medicine regimen incompletion. Reformulating the drug can result in delays in the development of a potential medicine, increasing the lead time to the patients. It might also require the use of extra animals in toxicity trials and lead to increased costs for pharma companies. We have developed a computational predictor for intense bitterness, that has above 80% accuracy. Applying the classifier to annotated datasets suggests that intense bitterness does not correlate with toxicity and hepatotoxicity of drugs. BitterIntense can be used in the early stages of drug development to identify drug candidates that require bitterness masking, and thus reduce animal use, time and monetary loss.

## Introduction

The use of sophisticated and highly automated processes in drug discovery have resulted in expedited pipelines and the approval of more than 4,000 medicines as of April 2020(1). Yet, a key aspect related to regimen compliance has not been properly addressed. The oral route remains the main way for drug administration(2), with aversive taste of drugs causing swallowing difficulties and compliance problems. This is especially relevant with pediatric medicine, for which encapsulation does not always provide a solution(3–5). Even though some bitter-masking agents exist, they are often insufficient for masking or preventing a drug’s intensely bitter taste(4–6). Indeed, more than 90% of pediatricians report that a drug’s aversive taste and low palatability are the biggest barriers to completing the medication regimen, leaving children with limited access to “child-friendly” drugs and exposing them to possible harm and insufficient treatment(4, 7). Because of the potential risks caused by aversive taste of drugs, the Food and Drug Administration (FDA) has added taste events to their Adverse Event Reporting System(8) and expects that all medicines with the potential to be given to children should be assessed for palatability(9, 10). The problem is also acute for the older population, and the European Medicines Agency (EMA) reflection paper on the development of medicines for geriatrics lists taste as a key consideration for medicine development(11). Sour or metallic taste can also elicit aversion, but intense bitterness is particularly abundant among drugs, and many of them were shown to activate bitter taste receptors(12, 13). The challenge imposed by aversive taste in drug development has led to the establishment of several assays for bitterness measurement, including the rat brief access taste aversion (BATA) (14), electronic tongues and human sensory panels(15).

The current pipeline for drug development process includes four main stages: drug design, preclinical phase (animal testing), clinical phases in human and the final review and approval by the medicines regulators including the FDA and the EMA(16, 17). During the design and preclinical phases, the taste of the drug is usually disregarded. It is evaluated, if at all, only during the clinical phases when the drug is introduced to humans. As a result, in clinical trials with nauseous drugs (usually due to intense bitterness), reduced compliance to the treatment and increased dropout rates have been documented in several cases(18, 19). In addition, knowledge of the aversive taste of drug candidates enables the selection of a similarly aversive placebo in order to avoid unblinding of the trials(20, 21). Often, it is only once intense bitter taste is suspected to affect the clinical trial and cause compliance problems, the palatability of the drug is tested, usually in human taste panels(15). This may lead to reformulation of the drug and repeat the preclinical and clinical phases. Such detours may delay a potential medicine from getting to the patient for an additional 6 to 24 months, potentially using additional animals, and increasing financial costs. For a moderately successful medicine, this could be a reduction in income estimated at hundreds of millions of dollars.

Bitter taste can be elicited by structurally diverse compounds. Over 1000 bitter-tasting compounds are currently documented in <u>BitterDB</u>. Salts, peptides, fatty acids, polyphenols, alkaloids, terpenoids and compounds from additional chemical families contribute to the wide chemical space of bitter tastants(22, 23). Bitter compounds vary also in their perceived intensity: quinine and amarogentin were reported as extremely bitter to humans, recognizable at micromolar concentrations. Other molecules, such as caffeine and naringin elicit slightly bitter taste and are typically recognizable at millimolar concentrations(23). Notably, we showed that bitter compounds are not more toxic than non-bitter compounds(24), questioning the common paradigm that posited the evolutionary role of bitter taste as a marker for toxicity(25, 26).

Bitter compounds are recognized by G-protein coupled receptors subfamily of bitter taste receptors, called T2Rs (27, 28), that harbor 25 functional T2R subtypes in human(29). While some receptors are broadly tuned with hundreds of diverse ligands, others are very selective with 0-3 known agonists(30, 31). T2Rs are not only expressed in the oral cavity but also in many extraoral tissues, possessing different physiological roles besides chemosensation of bitter tastants(32, 33). For example, activation of T2Rs expressed in human airway smooth muscle with inhaled bitter tastants was shown to mediate relaxation of the muscles and decreased airway obstruction in mouse models of asthma(34). T2Rs expressed in thyrocytes were shown to regulate the production of thyroid hormones and influence the function of the thyroid gland(35). Surprisingly, a human cohort suggested that polymorphism in T2R42 gene, an orphan bitter taste receptor, is associated with lower thyroid hormone levels(35). The fact that T2Rs can be expressed in extraoral tissues and that orphan T2Rs can be associated with physiological phenomena may suggest the involvement of T2Rs in health and disease(32) as well as a potential off-target for drugs(36).

We have previously developed BitterPredict(37), which classifies molecules into bitter or non-bitter with over 80% accuracy. Several other machine learning predictors followed suit(38, 39). In addition, structure-based methods were developed for identification of new agonists for specific bitter taste receptors(12, 40, 41). Here we are interested specifically in finding intensely or extremely bitter molecules, since these are the ones that are likely to cause compliance problems. This will allow project teams to address very bitter compounds in the early stages of development and focus on bitterness masking for the flagged molecules or deprioritizing them for oral route.

To establish a machine learning algorithm for intense bitterness, data had to be gathered: we curated data from BitterDB and from the literature, and have measured bitterness intensity of several new compounds using the BATA assay. Next, we successfully trained a new machine learning classifier “BitterIntense” with the ability to assign compounds as “very bitter” (VB) or “not very bitter” (NVB) based on descriptors calculated from their chemical structure. BitterIntense was then used to assess prevalence of very bitter compounds in datasets of interest in order to elucidate additional attributes of VB drugs.

## Results and Discussion

### Establishment of positive and negative sets

34 compounds were obtained from behavioral studies using the rat brief access taste aversion (BATA - see Experimental section). Additional compounds were pulled from the BitterDB(23), and the Analyticon repository of natural compounds on kaggle(42). The compounds were classified into 2 classes: “Very bitter” (VB) and “Not very bitter” (NVB) using the following criteria: Compounds with sensory bitter recognition threshold below 0.1mM; or molecules with taste description that states “extremely bitter” or “intensely bitter” etc., were included in the VB class (246 compounds). Compounds with bitter recognition threshold above 0.1 mM or molecules with taste description that includes “slightly bitter” or “weak bitter taste” etc., were included in the NVB class (323 compounds). The BATA test measures aversion, which is assumed to be driven mainly by bitterness. The IC_50_ achieved for each compound in the BATA test was used to classify compounds using the following criteria: molecules with IC_50_ below or equal to 3mM were classified as VB. Molecules with IC_50_ above 3mM were classified as NVB. In addition, 152 non-bitter compounds were added to the NVB class from the negative set used previously in BitterPredict(37) since for practical purposes pursued here, non-bitter and not very bitter fall under the same category. The non-bitter compounds that were included in the negative set were randomly selected and had MW>250 g/mol to match the MW of very bitter compounds.

### Chemical families analysis

The chemical families for VB and NVB compounds in the training set are represented in Figure 1. The VB class is enriched with triterpene saponins and triterpenes in general, while NVB class is broadly represented by different chemical families.

**Figure 1.**
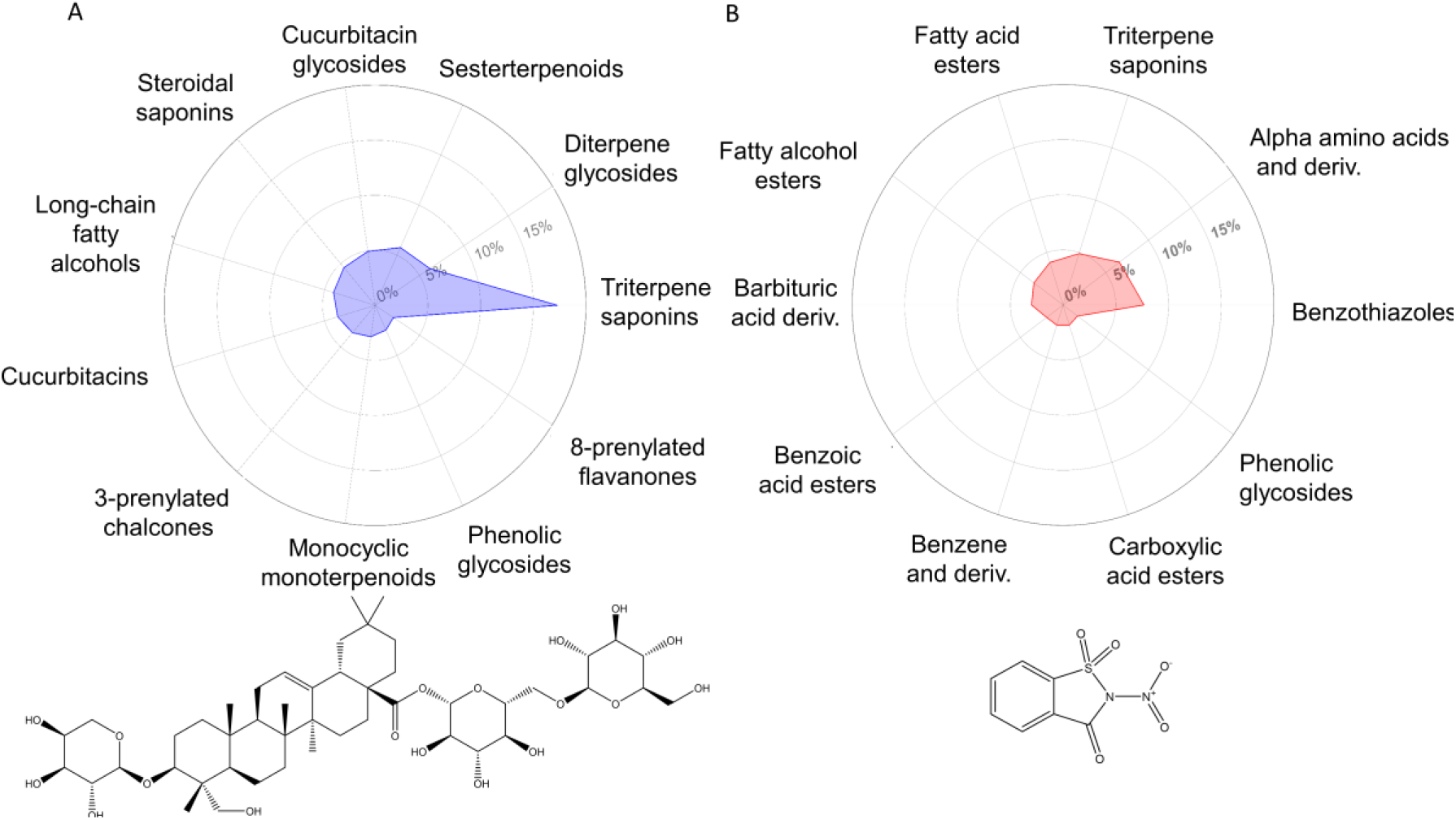
Representation of chemical families in the training set (A) – Top chemical families represented in the dataset of very bitter compounds. (B) - Top chemical families represented in the dataset of not very bitter compounds. The compound on the lower left side (Asperosaponin VI) is a representative of very bitter triterprene saponins, the compound on the lower right (Nitrosaccharin) is a representative of not very bitter benzothiazoles. Deriv. = derivatives

### Training the classifier

Physicochemical, Ligfilter and Qikprop descriptors were calculated for the compounds in the dataset (see Experimental Section). 15% (105) of the compounds were chosen randomly and left out of the training set, as a hold-out test set for final evaluation of the model. The other 616 compounds were randomly divided into: 80% training set (493 compounds) and 20% internal test set (123 compounds). Extreme Gradient Boosting (XGBoost) algorithm was chosen for this classification task. XGBoost is a popular and powerful decision-tree-based ensemble method that uses optimized gradient boosting techniques and is known to perform well with small to medium size datasets(43).

Since the training set is rather small, we extracted the highest contributing features (see Experimental Section) in order to avoid overfitting. 55 features were selected out of 235 for the model training. Further details are described in the Methods section.

### BitterIntense Performance

Evaluation of the model’s performance (Table 1) was carried out on three sets: Training set (with cross-validation, k-fold = 10), Test set, and Hold-out set. BitterIntense was able to achieve over 80% accuracy across the different datasets. In general, we observed higher recall values than precision values. This result is in line with our goal to maximize identification of very bitter compounds.

**Table 1:** BitterIntense performance on the training, test and hold-out sets. Training set evaluation was done using k-fold cross validation with k =10. The results in the training set column represent the mean metric with its standard deviation across 10 iteration of cross validation.

|  | Training set<br>(493<br>compounds) | Test set<br>(123<br>compounds) | Hold-out set<br>(105<br>compounds) |
| --- | --- | --- | --- |
| Accuracy (%) | 87±5 | 83 | 80 |
| Precision (%) | 80±8 | 71 | 63 |
| Recall (%) | 85±4 | 86 | 77 |
| Specificity (%) |  | 81 | 81 |

### Important features

The feature importance was measured in XGBoost by the “gain” method which is the average gain of splits which use the feature in the prediction process. Higher gain value implies greater importance of the feature. The top 15% of features are represented in Figure 2 and suggest that molecule’s size and polarizability are the most influential factors for bitterness level. Fitting the model using the heavy atom count feature only, results in 70% accuracy on the training set, with recall and precision of 70% and 58% respectively. This means that VB molecules are often larger than NVB molecules, but additional features greatly improve the predictions, reaching average accuracy of 87%, recall of 85% and precision of 80% on training set. An important contributing feature is molar refractivity, which is a measure of a compound’s polarizability.

**Figure 2.**
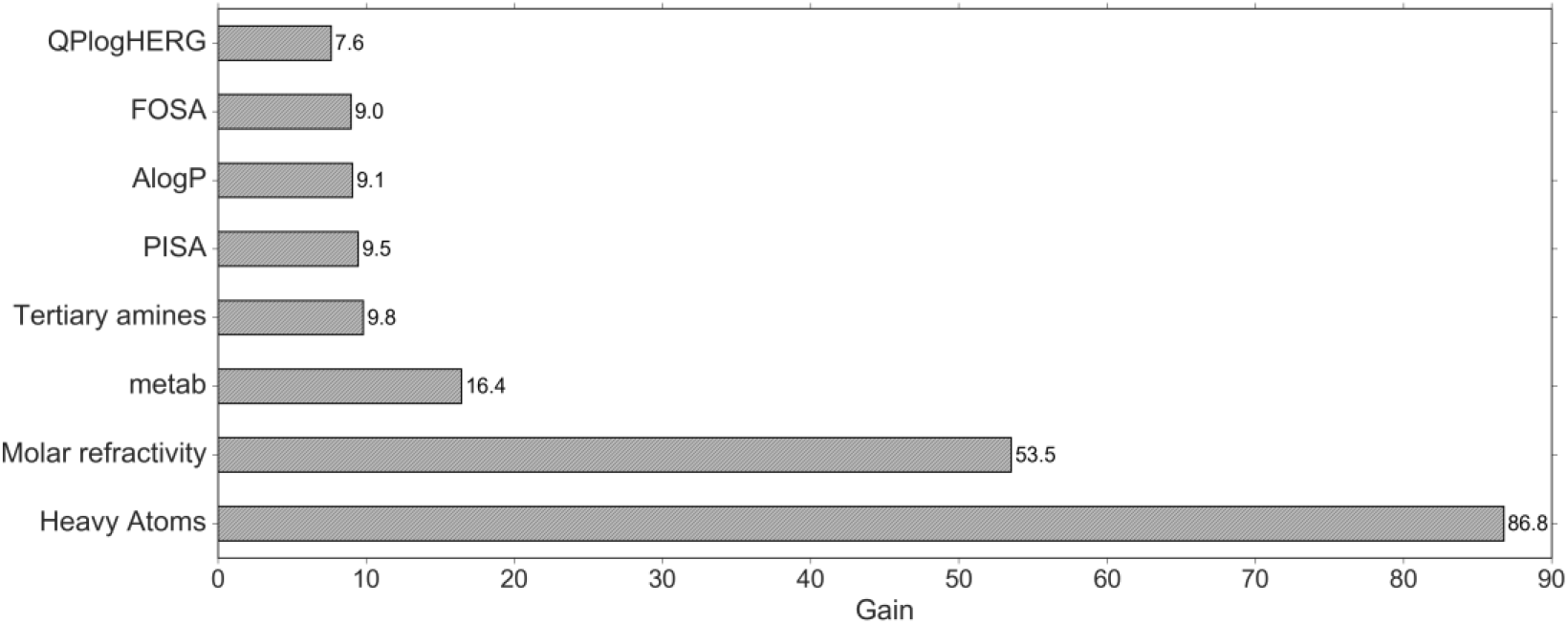
Top 15% important features in the model. The importance is calculated by the average gain of splits which use the feature in the prediction process in each tree in the XGBoost model. The heavy atom count, molar refractivity, number of likely metabolic reactions (metab), number of tertiary amines and amide groups, π (carbon and attached hydrogen) component of the solvent accessible surface area (PISA), hydrophobicity (AlogP), hydrophobic component of the solvent accessible surface area (FOSA) and Predicted IC50 value for blockage of HERG K+ channels (QPlogHERG).

### Relationship between bitterness intensity and toxicity

Though bitter taste is often regarded as a marker for toxicity that guards against consuming poisons(44), our previous analysis showed that bitter compounds are not necessarily toxic and vice versa(45). Here we apply BitterIntense to the FocTox and CombiTox datasets (45). The FocTox dataset consists of FAO/WHO food contaminants and extremely hazardous substances.

Out of 289 compounds, only 25 were predicted to be very bitter (pVB, 8.6%). CombiTox, a manually curated dataset of toxic compounds, was also analyzed for very bitter substances. Out of 134,057 compounds, 12% were pVB, suggesting that toxic substances are not necessarily very bitter and in fact, most of the toxicants are predicted to be NVB.

We further checked the possible connection between the level of bitterness and liver toxicity (hepatotoxicity). Hepatotoxicity is the most common cause for the discontinuation of clinical trials on a drug, and the most common reason for an approved drug’s withdrawal from the marketplace(46, 47). Drug-induced liver injury (DILI) has been listed as the leading cause of acute liver failure in the USA in 2002(48), and DILI has become an important concern in the drug discovery process. A possible connection between the level of bitterness and hepatotoxicity could suggest that the level of bitterness is a potential marker for such toxicity. We predicted the level of bitterness of drugs with known hepatotoxicity descriptors, taken from DILIrank dataset(46).

Most of the drugs in the dataset (729 compounds) were predicted as NVB (pNVB), and only 258 were predicted as VB. The pVB drugs do not appear to be more hepatotoxic than the pNVB drugs: the most hepatotoxic class (class number 8, Figure 3A) as well as the “Most DILI concern” category (Figure 3B) are actually enriched with pNVB drugs There are several possible explanations for this trend: some bitter compounds that activate T2Rs were also shown to interact with Cytochrome P450 (CYP)(49), promiscuous monooxygenase enzymes involved in phase 1 metabolism of drugs and xenobiotics in the human body(50). Perhaps VB and pVB drugs interact differently or even with stronger affinity with CYP enzymes, possibly explaining the differences in the hepatotoxic effect. Furthermore, activation of extraoral T2Rs might contribute to the decreased risk for hepatotoxic effect of the very bitter drugs. It was previously shown that activation of gut T2Rs had let to detoxification effect by upregulating the transcription of xenobiotic efflux pumps(51). Such effects might also take place in the liver in addition to the gut. From these results we can conclude that the preference for pNVB drug candidates over pVB drugs are not related toxicity considerations. In fact, our analysis shows that in many cases the VB and pVB drugs tend to be less harmful and should not be automatically discontinued in the drug discovery process.

**Figure 3.**
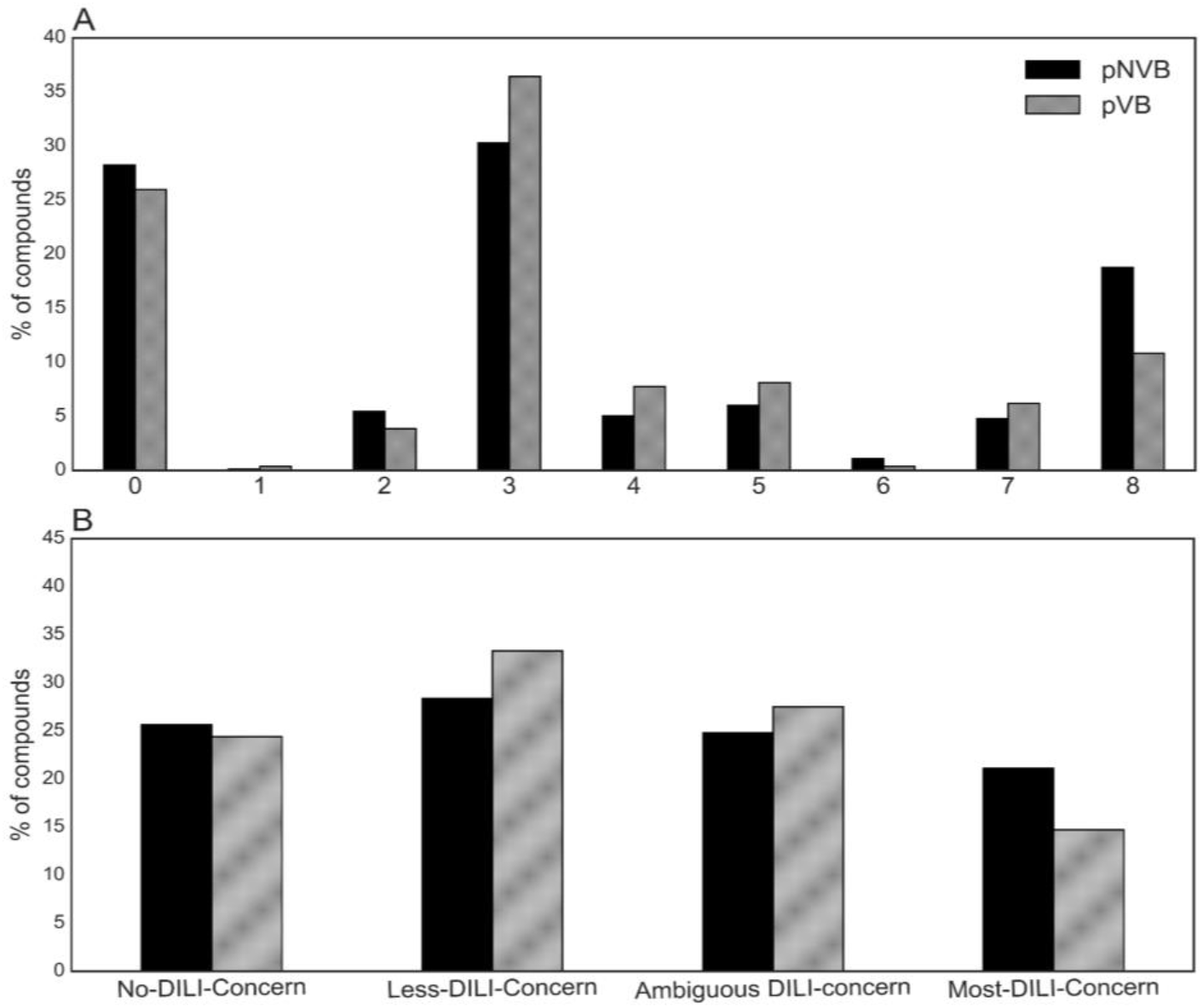
Bitterness levels of drugs and their hepatotoxicity descriptors. (A) Distribution of pVB (silver) and pNVB (black) drugs across severity classes of hepatotoxicity. (B) Distribution of pVB and pNVB drugs across DILI concern categories. The severity of hepatotoxicity increases from left to right in all figures.

### Very bitter drugs and their potential therapeutic effects

Since VB drugs are more likely to cause compliance problems, we applied BitterIntense on all compounds (approved and experimental drugs) from Drugbank (5.1.5)(1) to evaluate the abundance of VB drugs (figure 4). Out of 10,170 compounds that were able to pass through our predictor, 23.6% were pVB. Specifically, 18% of experimental drugs and 26% of approved drugs are pVB. For comparison, in microbial natural products (NPatlas, version 2019_08, n=24,805) (52), 47.7% were pVB. Thus only ¼ of drug candidates, but about half of the microbial natural compounds are likely to be VB.

**Figure 4.**
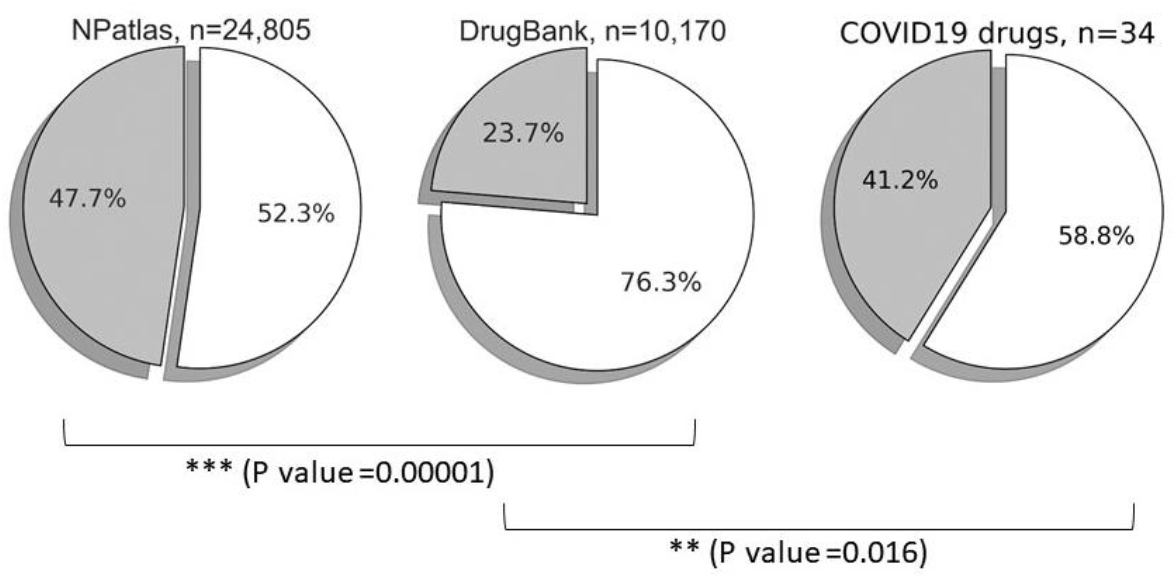
Prevalence of pVB compounds (silver) and pNVB compounds (white) across 3 datasets: NPatlas, DrugBank and COVID19 drugs. Statistical significant difference in the proportions of pVB compounds was observed using Two Proportion Z-Test.

Are there specific therapeutic indications or targets that are enriched with pVB drugs? We focused here on a highly relevant disease, the COVID19. The COVID19 pandemic is spreading throughout the world, with millions of confirmed cases and hundreds of thousands of deaths, according to reports by the World Health Organization that were published in June 2020(53). Some drugs have been suggested for treating COVID-19 patients, but no drug has yet been approved fully and officially by the FDA. Several drugs are currently under study and clinical trial, for example: Remdesivir, an adenosine analog that was previously tested as a potential drug for Ebola and as anti-viral drug(54), showed promising result in COVID19 patients and thus was granted an FDA Emergency Use Authorization on 1 May 2020(55). Interestingly, taste and smell loss,(56) including impairment of the bitter taste, are reported by many COVID19 patients(57). We applied BitterIntense to possible COVID19 drug candidates and compared it to the general abundance of pVB drugs in DrugBank. A list of ligands related to COVID19 in “Coronavirus Information - IUPHAR/BPS Guide to Pharmacology”(58) was retrieved. After excluding antibodies and compounds without chemical structure (Figure 4), among 34 drug candidates, 41.2% were pVB. The proportion of pVB drugs among COVID19 potential drugs, is thus significantly higher than in DrugBank (23.6%)(P-value = 0.016), suggesting that VB drugs may be more abundant in the COVID19-related list than in general drugs. No evident difference was found between the main targets of pVB and NVB COVID19-related drug candidates. While the potential involvement or mediation of bitter taste receptors in COVID19 is unclear, the results further highlight the importance of flagging - but not excluding from the pipeline – of pVB drugs and drug candidates.

In conclusion, BitterIntense was developed to easily classify compounds as Very bitter or Not very bitter and has achieved above 80% accuracy on test set. This quick and easy method enables the identification of intensely bitter compounds during early stages of drug development, potentially reducing financial costs, animal testing and the lead time to the patients The ability to detect pVB drugs in early stages not only will accelerate the drug development process but will also promote the development of more palatable drugs suitable for children and geriatric patients. However, intensely bitter molecules should not be eliminated completely from the development process. Our analysis revealed that such compounds are not necessarily more toxic than pNVB compounds, and not more hepatotoxic. Furthermore, we found that potential COVID19 drugs were enriched in intensely bitter molecules, particularly interesting in view of possible involvement of taste impairment in COVID19 disease(59, 60).

## Materials and Methods

### BATA assay

The rat brief access taste aversion (BATA) model has been demonstrated as a highly translatable tool for screening bitter compounds(14, 61). Comparison of BATA and human gustatory trials at GlaxoSmithKline (GSK) suggest an average offset of 0.5 log concentration, with rats typically slightly more tolerant of bitter taste than humans(61). Studies at GSK were performed using Davis Rig MS-160 lickometers (DiLog Instruments, Tallahassee, USA) and as described by Soto et al(14) with the following exceptions. 12 male Sprague Dawley Crl:CD (SD) rats (number determined following power analysis of historic GSK data), 6 to 7 weeks of age on arrival, supplied by Charles River UK (Marston, UK) were used per study. Rats were housed in groups of either two or four, kept on a 12 hour light: dark cycle, 19-21oC, 45-55% humidity. 5LF2 rodent diet (LabDiet, Missouri, USA) was fed ad libitum. Animal grade drinking water (AGW) was reverse osmosis filtered, UV treated and provided ad libitum between water restriction periods. All testing occurred during the light period. Rats were water restricted for 21 hours prior to each test session to ensure sufficient thirst for the rats to attempt to lick all solutions presented. Each test session was limited to a maximum of 30 minutes, following which rats were returned to their home cages and given free access to AGW for a minimum of 2.5 hours before commencement of the next water restriction period. Following completion of each study, rats were health assessed by a named veterinary surgeon and returned to non-naïve stock. A minimum one-week washout period was provided prior to use of rats on subsequent BATA studies.

All compounds presented during BATA studies were fully solubilised across a range of concentrations with 0.5 log dose separation to generate concentration response curves. Lick counts of zero and one were excluded from data sets due to being deemed an insufficient attempt to lick a solution and therefore not related to palatability. Lick responses were modelled using a three parameter logistic function with the minimum constrained to zero (R v.3.5.1, Foundation for Statistical Computing, Vienna, Austria). Lick response was expressed as a percentage of the median AGW response within each study, 95% confidence interval. The concentration of API that elicited lick rates equivalent to 50% of the median AGW was then calculated and deemed the IC50.

All animal work conforms to the UK Animals (Scientific Procedures) Act, European Directive 2010/63/EU and the GSK Policy on the Care, Welfare and Treatment of Animals. All protocols were approved as part of the GSK Scientific and Ethical Review Forum and carried out in accordance with the appropriate project licence issued by the UK Home Office.

For all studies performed at GSK, commercially available compounds were sourced from Sigma Aldrich (Gillingham, UK). GSK compounds were supplied by the internal dispensary.

### BitterDB and Analyticon data

Data on bitterness intensity and chemical structures for the training and testing of the model were also obtained from BitterDB(23) and Analyticon’s repository on Kaggle website(42).

The addition of the non-bitter subset had shown to be beneficial in preliminary testing of the classifier (not shown), increasing the ability to distinguish between very bitter compounds and not very bitter compounds.

### Chemical families analysis

The chemical families of the compounds in the training set were extracted using ClassyFire webserver(62).

### Datasets preparation

After obtaining the SMILES strings of the compounds, we uploaded the compounds to Maestro (Schrödinger Release 2017–2: MS Jaguar, Schrödinger, LLC, New York, NY, 2017). We generated 3D structures using ligprep and Epik (Schrödinger Release 2017-2: LigPrep, Epik, LLC, New York, NY, 2018) in physiological pH 7.0 ± 0.5. All compounds were desalted when available and we retained the original chirality of compounds when specified, otherwise all stereoisomers were generated. For each compound, the conformer with the lowest energy was extracted and used. When 2 protomers where generated for one compound, we kept both structures. Compounds that could not be neutralized were excluded from the sets due to the limitations of calculating QikProp descriptors. All the datasets in this current study were prepared in the same protocol as mentioned above.

### Descriptors calculation

Three sets of descriptors were calculated for the prepared 3D structures using Canvas (Schrödinger Release 2017-4: Canvas, Schrödinger, LLC, New York, NY, 2017): Physicochemical descriptors, Ligfilter descriptors (moieties, atoms and functional groups) and QikProp descriptors (ADME descriptors). For the QikProp descriptors, additional PM3 properties were calculated as well (Schrödinger Release 2017-4: QikProp, Schrödinger, LLC, New York, NY, 2017). Compounds that failed to calculate one of the descriptors were excluded from the analysis.

### Model construction and fitting

The XGBoost model was constructed and fitted using Python 3.7.5, in Spyder 3.7 environment. Relevant packages: Scikit-learn (version 0.21.3), XGBoost (version 0.9), Numpy (version 1.17.4), pandas (version 0.25.3), matplotlib (version 3.1.1) and seaborn (version 0.9.0).

An early stopping approach was used in the training process, in which the performance of a model is monitored during the training process which is stopped once the performance ceased improving, see Figures S1 and S2. The evaluation metrics for the early stopping were: logarithmic loss(63) and Binary classification error rate (defined as = #(wrong cases)/#(all cases)).

### Feature selection

The most contributing features were selected according to their feature importance gain score, calculated using XGBoost library in python(43). We constructed a loop that tested the changes in the accuracy of the model by setting different thresholds of the gain scores in order to select the best features. When setting the threshold on 0.004, we maintained 55 features (out of 235 original features) that had the most contribution to model, obtaining 83% accuracy.

### Parameter tuning

The parameters of the XGBoost algorithm were tuned using sklearn’s GridSearchCV from sklearn.model selection module(64). The number of cross validation folds was set to 10 and the scoring method was set for ‘f1 score’ in order to improve the precision and the recall. Parameter tuning was performed on the training set using the initial fitted model, suggesting that the optima parameters are: colsample_bytree=0.6, gamma=0.5, max_depth=5. In brief, colsample_bytree is the fraction of features that will be randomly sampled to construct each decision tree, gamma represents the minimum loss reduction required to make a further partition on a leaf node of the decision tree and max depth represents the maximum depth of a tree. In addition, the parameter (scale_pos_weight) that helps with unbalanced data was tuned by the ratio between positive and negative observations in the training set to 1.9.

### Evaluation of the performance of the model

After calculating the number of true positives (TP), true negatives (TN), false positives (FP) and false negatives (FN), we evaluated the model using four metrics:

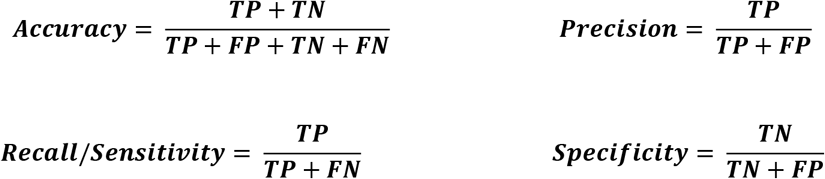

### Toxicity data

FocTox dataset consists of FAO/WHO food contaminants list and a list of extremely hazardous substances defined in section 302 of the U.S. Emergency Planning and Community Right-to-Know Act. CombiTox dataset is a combination of two datasets (The Toxin and Toxin-Target Database version 2.0 (T3DB) and DSSTox—the Distributed Structure-Searchable Toxicity Database). The datasets were taken from the paper of *Nissim I. et al*(45).

### Hepatotoxicity data

All hepatotoxicity descriptors were extracted from FDA’s DILIrank(46) dataset which is an updated version of the LTKB (Liver Toxicity Knowledge Base) Benchmark dataset(65). DILIrank consists of 1,036 FDA-approved drugs with known hepatotoxicity descriptors and liver toxicity risk assessments. The compounds in the dataset were prepared as explained in “Datasets preparation” section.

### External datasets

DrugBank (version 5.1.5) consist of experimental and approved drugs(1), and Natural products atlas (NPatlas, version 2019_08)(52) were downloaded from their official websites. Compounds were prepared according to the protocol of “Datasets preparation” section. COVID 19 drugs and their targets were retrieved from “IUPHAR/BPS Guide to Pharmacology”(58). After excluding the antibodies and compounds without chemical structure we prepared the remaining 34 compounds according to the “Datasets preparation” section.

### Analysis and visualization of the data

All the data in this current study was analyzed using Pandas library(66) and visualized with Matplotlib(67) library in Python.

## Acknowledgments

MYN is grateful to the late Professor Amirav Gordon for his invaluable advice, encouragement and support. The authors thank Yuli Slavutsky and Dr. Yuval Benjamini for helpful discussions on machine learning methods. The core data science scholarship to EM from the Center for Interdisciplinary Data Science Research (CIDR) is gratefully acknowledged. MYN and EM participate in Mu.Ta.Lig (CA15135) and ERNEST (CA18133) COST actions. MYN is supported by the Israel Science Foundation grants ISF 494/16 and ISF-NSFC2463/16.

