## Supplementary figures for "Intense bitterness of molecules: machine learning for expediting drug discovery"

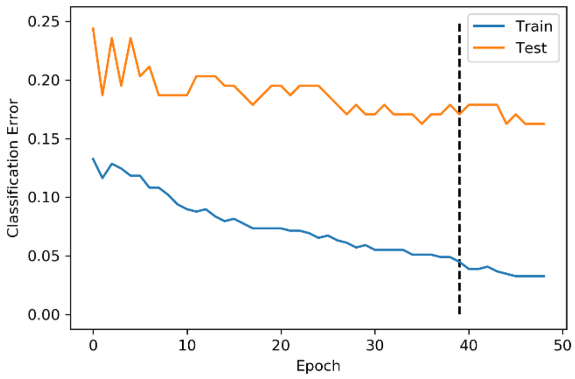


**Figure S1**. Classification error rate (CER) throughout the training epochs during the early stopping training process. Each epoch one training iteration. In epoch number 39 (dashed line) the CER in the training set keeps decreasing while in the test set the CER increasing, suggesting this epoch is the as a potential stopping point.

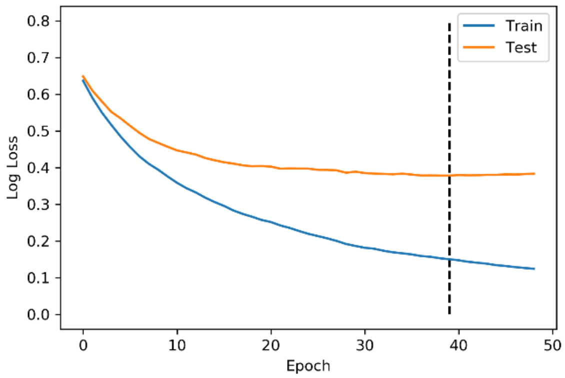


**Figure S2**. Logarithmic loss (log loss) throughout the training epochs during the early stopping training process. Each epoch represents one training iteration. In epoch number 39 (dashed line) the log loss in the training set keeps decreasing while in the test set the log loss remains steady.
